# Fox: Predicting rates of host-symbiont co-evolution events with association-aware attention

**DOI:** 10.64898/2026.10.29.762770

**Authors:** Gabriele Di Palma, Laurent Jacob, Catherine Matias, Blerina Sinaimeri

## Abstract

**Motivation:** Hosts and their symbionts share a history of associations in which lineages cospeciate, switch hosts, duplicate, or are lost. Quantifying the relative frequencies of these four events is a central goal of cophylogenetics, but existing tools typically require a fully resolved, correctly rooted host tree, often alongside a symbiont tree, propagating tree-inference errors into the reconciliation and scaling poorly to large systems.

**Results:** We introduce Fox, a transformer neural network architecture that infers the relative frequencies of the four cophylogenetic events directly from a pair of host and symbiont multiple sequence alignments (MSAs) and their sequence mapping, with no tree-reconstruction or reconciliation step. Fox is trained under a simulation-based inference scheme: it learns event frequencies from alignments generated by a coevolutionary process, paying its computational cost once at training time and processing any new system in a single forward pass. On held-out data from its training simulator, Fox recovers event frequencies accurately and with near-zero bias. On additional synthetic data from an independent cophylogeny simulator, its error roughly doubled, but is still very low allowing to track the true frequencies. Most importantly, when compared to the Approximate Bayesian Computation (ABC)-based method AmoCoala on these data, it reduces mean absolute error by 45-76% for cospeciations, host switches and duplications and shows similar error for losses, while running more than three orders of magnitude faster. Applied to the empirical cockroach-Blattabacterium system, Fox recovers the expected cospeciation-dominated profile, showing that behaviour learned on simulated data transfers to real sequences. These results confirm that the coevolutionary signal usually extracted from trees can be read directly, accurately, and fast in the alignments themselves.

**Availability:** Fox can be downloaded from https://github.com/Gabbo240900/Fox

**Key Messages:**

- Fox infers the relative frequencies of the four cophylogenetic events directly from paired host and symbiont MSAs and their mapping, with no tree-reconstruction step.
- Fox produces inferences faster than AmoCoala and is able to scale to large symbiont trees that AmoCoala could not complete.
- On simulated data from an independent simulator, Fox is more accurate and less biased than the ABC baseline AmoCoala, and on empirical data it recovers the expected cospeciation-dominated profile.

## Introduction

Hosts and their symbionts are connected by ecological associations that may persist over long evolutionary periods. As host species diversify, their associated symbionts may undergo different evolutionary processes: they may diversify in correspondence with their hosts, colonize new host species, diversify independently within the same host, or fail to persist in some descendant hosts. The co-evolutionary patterns observed today therefore reflect the combined action of several processes acting on the two interacting set of species and on their associations (Braga et al. 2018; de Vienne et al. 2013; Dismukes et al. 2022).

Understanding the relative importance of these processes is a central question in cophylogeny (Page 1994; Dismukes et al. 2022; Charleston 2003). Different evolutionary events correspond to substantially different biological scenarios. *Cospeciation* occurs when the symbiont diverges in correspondence to the divergence of a host species; *host switching*, in contrast, occurs when a symbiont switches from one host species to another independently of any host divergence; *duplication* happens when the symbiont diverges but not the host; while *loss* represents the absence, in one of the descendant hosts, of a symbiont association expected from the reconstructed evolutionary history (Page 1994; Baudet et al. 2014). Importantly, an inferred loss does not necessarily correspond to a directly observed extinction: depending on the biological system and the available data, it may also reflect failure of the symbiont to persist after host speciation or failure to observe or sample an existing association (Dismukes et al. 2022).

Distinguishing among these processes is biologically important, as similar host-symbiont phylogenetic patterns can arise from different evolutionary mechanisms. In particular, phylogenetic congruence does not necessarily imply cospeciation, since host switching followed by diversification can produce similar patterns (Vienne et al. 2007). Estimating the contributions of cospeciation, host switching, duplication, and loss can thus clarify host associations and diversification, particularly in host-pathogen systems (Parrish et al. 2008; Longdon et al. 2014), but also in parasites, endosymbionts, and mutualists (Dismukes et al. 2022).

Computational approaches to cophylogenetics include global-fit methods, such as ParaFit (Legendre et al. 2002) and PACo (Balbuena et al. 2013), which test for overall correspondence between host and symbiont phylogenies but do not identify the evolutionary events generating it. *Event-based reconciliation methods* instead provide an explicit evolutionary explanation by mapping a symbiont phylogeny onto a host phylogeny through events such as cospeciation, duplication, host switch, and loss (Page 1994; Charleston 1998). Classical approaches assign costs to these events and search for reconciliations minimizing the total cost (Charleston 1998; Conow et al. 2010). This formulation has two important limitations. First, the inferred evolutionary scenario can depend strongly on the event costs supplied to the algorithm: different cost combinations may favour substantially different reconciliations (Huber et al. 2018; Santichaivekin et al. 2019; Wang et al. 2020, 2022). Moreover, several distinct reconciliations may achieve the same minimum cost, so that a single optimal score does not necessarily identify a unique biological history. Second, reconciliation can become computationally demanding when host-switch events and temporal consistency constraints are incorporated. In particular, finding optimal time-consistent reconciliations on undated host trees is computationally difficult in general, motivating the use of heuristics, approximations, or additional assumptions in practical reconciliation methods (Conow et al. 2010).

These limitations motivated statistical approaches for estimating evolutionary-event parameters. Coala (Baudet et al. 2014) uses Approximate Bayesian Computation (ABC) to infer cospeciation, duplication, host-switch, and loss probabilities, while AmoCoala (Sinaimeri et al. 2023) extends this framework to multiple host-symbiont associations. The method of Alcala et al. (2017) also uses ABC, but focuses on cospeciation and host-switch rates while accounting for phylogenetic uncertainty through multiple dated symbiont trees. A related reconciliation problem arises in the gene-species setting, where similar event-based models are used to describe gene evolution within species histories. In this context, ALE (Szöllősi et al. 2013) uses a likelihood-based framework to infer duplication, transfer, and loss while accounting for gene-tree uncertainty.

Despite their advantages, these methods share two main limitations:

- **Dependence on phylogenetic reconstruction**. A host phylogeny, and in some cases also a symbiont phylogeny, is required as input. Errors or uncertainty in these phylogenies can affect the subsequent estimation of coevolutionary events.
- **High computational cost**. Inference can become computationally demanding for large systems. ABC-based approaches require repeated simulations for each new dataset, while likelihood-based methods must evaluate many possible reconciled histories.

These limitations motivate a different formulation of the problem: inferring coevolutionary event frequencies directly from host and symbiont multiple sequence alignments (MSAs) and their association mapping, without reconstructing the corresponding phylogenetic trees. We address this problem by training a deep learning model on simulated host-symbiont systems for which the true event frequencies are known. The simulations are therefore required only during training; once trained, the model can analyse a new host-symbiont system directly from its MSAs and associations, without additional simulations or tree reconstruction.

This approach builds on recent advances in neural methods for evolutionary inference. Previous work has shown that likelihood-free inference methods exploiting neural networks could estimate phylogenies (Nesterenko et al. 2025; Blassel et al. 2026) or related evolutionary parameters such as birth-death rates (Garot et al. 2026) directly from MSAs. These precedents suggest that a neural inference approach for cophylogeny event rates from MSAs is feasible. Nonetheless, the challenge is to design a neural network architecture that jointly attends two alignments (host and symbiont) and accounts for the mapping that links the taxa of the two organisms.

We introduce Fox, a transformer-based neural network that outputs the relative frequencies of the four cophylogenetic events directly from a pair of MSAs and a host-symbiont sequence mapping with no tree inference. Fox combines axial attention over each alignment with a pair-bias track and association-aware attention, in which each host–symbiont link becomes a token, so that the evolutionary signal can be shared between the two lineages. The network trained on data simulated under a host-symbiont co-evolution model can be proved to formally approximate the posterior distribution of the rates conditional to the MSAs (see *e*.*g*., Radev et al. 2020; Blassel et al. 2026) or quantiles of this distribution (Garot et al. 2026).

We use AsymmeTree (Schaller et al. 2022) to generate coevolutionary histories with known event frequencies and the corresponding host and symbiont phylogenies, which are then used to simulate molecular sequences. We first evaluate Fox on held-out datasets generated by AsymmeTree to assess its accuracy under the training simulation model. To study its ability to generalize beyond this model, we also evaluate Fox on datasets generated by the independent simulator treeducken (Dismukes and Heath 2021). On these datasets, we compare Fox with AmoCoala, a tree-based ABC method that estimates cophylogenetic event probabilities through simulation-based inference. Finally, we apply Fox to an empirical host-symbiont dataset as a proof of concept.

On held-out AsymmeTree data, Fox recovers event frequencies with low errors and nearly zero bias, achieving high *R*^2^ (variance explained) for host switches and duplications, and moderate performance for cospeciation and losses. On treeducken data, its errors roughly double, but predictions remain correlated with the true frequencies, with positive *R*^2^ for cospeciation, host switches, and duplications, and a slightly negative *R*^2^ for losses. Compared with AmoCoala on the same treeducken datasets, Fox achieves errors two to four times smaller for cospeciation, host switches, and duplications, while performing similarly for losses. On an empirical dataset, both methods recover the expected cospeciation-dominated pattern, while Fox requires substantially less computation at inference time. These results show that, once trained, deep learning models can rapidly estimate cophylogenetic event frequencies directly from sequence alignments, shifting the computational cost to a one-time training phase and avoiding additional simulations or phylogenetic reconstruction for each new dataset.

Our contributions are:

- A new formulation of cophylogenetic event-frequency estimation as a supervised learning problem from paired host and symbiont MSAs and their association mapping, avoiding phylogenetic tree reconstruction at inference time;
- Fox, an asymmetric dual-MSA transformer specifically designed to model host–symbiont evolution through association-aware attention;
- An evaluation of Fox on simulated and real data, showing higher accuracy than AmoCoala on simulated datasets with known event frequencies and substantially faster inference once the model is trained, together with a proof-of-concept application to an empirical host-symbiont system.

## Results

The experiments conducted with Fox were driven by three main questions:

- Is Fox able to learn from simulated data how to estimate event-frequencies from host/symbiont MSAs and their associations?
- Does Fox perform better than other models at predicting event frequencies in a misspecified setting on data simulated under a model different from the one used to learn our estimator?
- Given some real co-phylogenetic data, is Fox able to predict event frequencies and how does it compare to other methods?

### Validation of event-frequency estimation

The first step was to assess whether Fox was able to correctly learn event frequencies on AsymmeTree synthetic data generated following the setup defined in the *Tree and sequences simulation* subsection.

The model has been evaluated on the validation set using six complementary metrics per event: mean absolute error (MAE) and root-mean-squared-error (RMSE), variance explained (*R*^2^) and linear correlation with the truth (Pearson *r*), mean-relative error (MRE, that is informative for rare events whose absolute frequencies are small), and signed bias (mean prediction minus truth, capturing systematic over or under estimation). Fox recovers all four event frequencies closely (Table 1), with *R*^2^ values of 0.78, 0.85, 0.60 and 0.83 for cospeciation, host switch, loss and duplication, respectively. Its average MAE across all four events is 0.028 and it shows almost no bias (|bias| *≤* 0.005). From training and validation, we could assess that loss is the hardest event to predict.

**Table 1.** Validation metrics of Fox on AsymmeTree-simulated data.

| | MAE | RMSE | $R^2$ | Pearson $r$ | MRE | bias |
| --- | --- | --- | --- | --- | --- | --- |
| Cospeciation | 0.0441 | 0.0578 | 0.777 | 0.883 | 0.072 | +0.005 |
| Host-Switch | 0.0163 | 0.0220 | 0.848 | 0.925 | 0.164 | -0.005 |
| Loss | 0.0296 | 0.0387 | 0.597 | 0.776 | 0.268 | -0.004 |
| Duplication | 0.0204 | 0.0261 | 0.829 | 0.913 | 0.117 | +0.004 |

Figure 1 shows per-event scatter plots on training and validation sets. Predictions concentrate on the identity line for all four events, with the largest residual variance on losses, consistent with its smaller marginal support in the simulator. In Supplementary Figure 3, we also show the 2-D density plots for the same training and validation sets.

**Figure 1.**
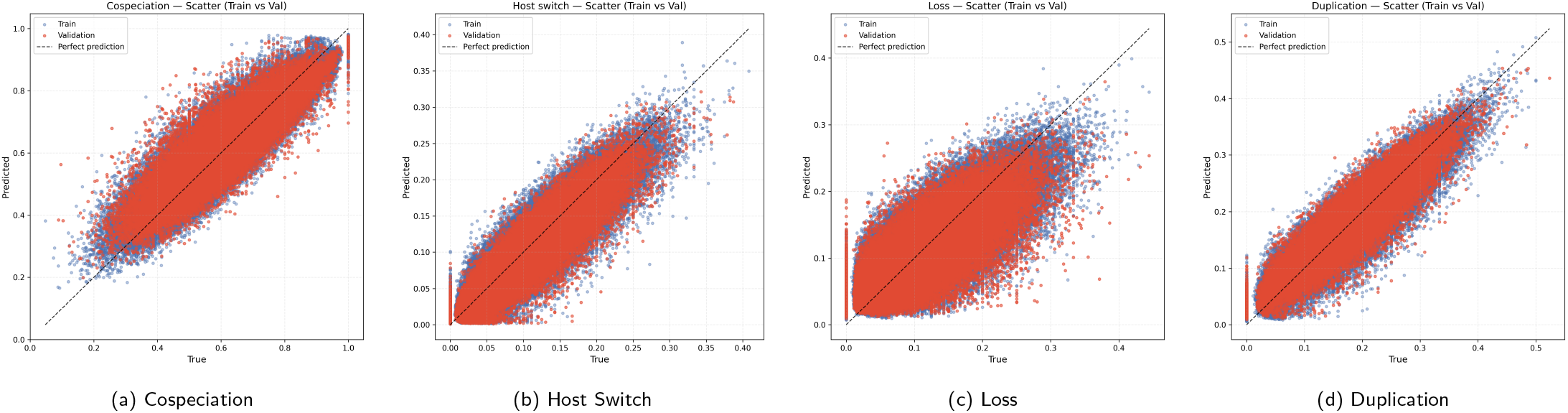
Predicted vs. true event frequencies on train and validation set. The dashed line is the identity.

**Figure 2.**
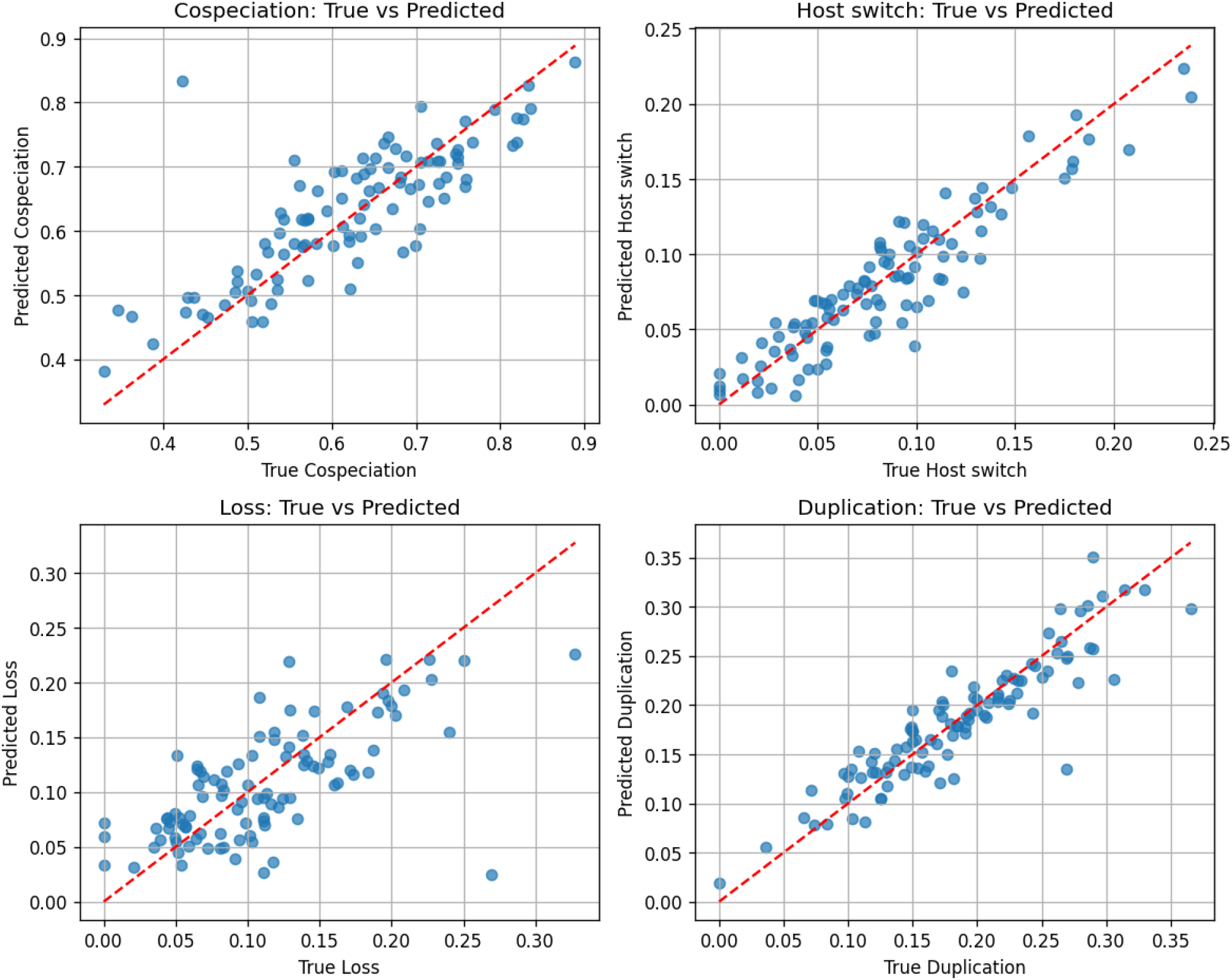
Fox predictions on the AsymmeTree held-out test set (*n* = 100).

**Figure 3.**
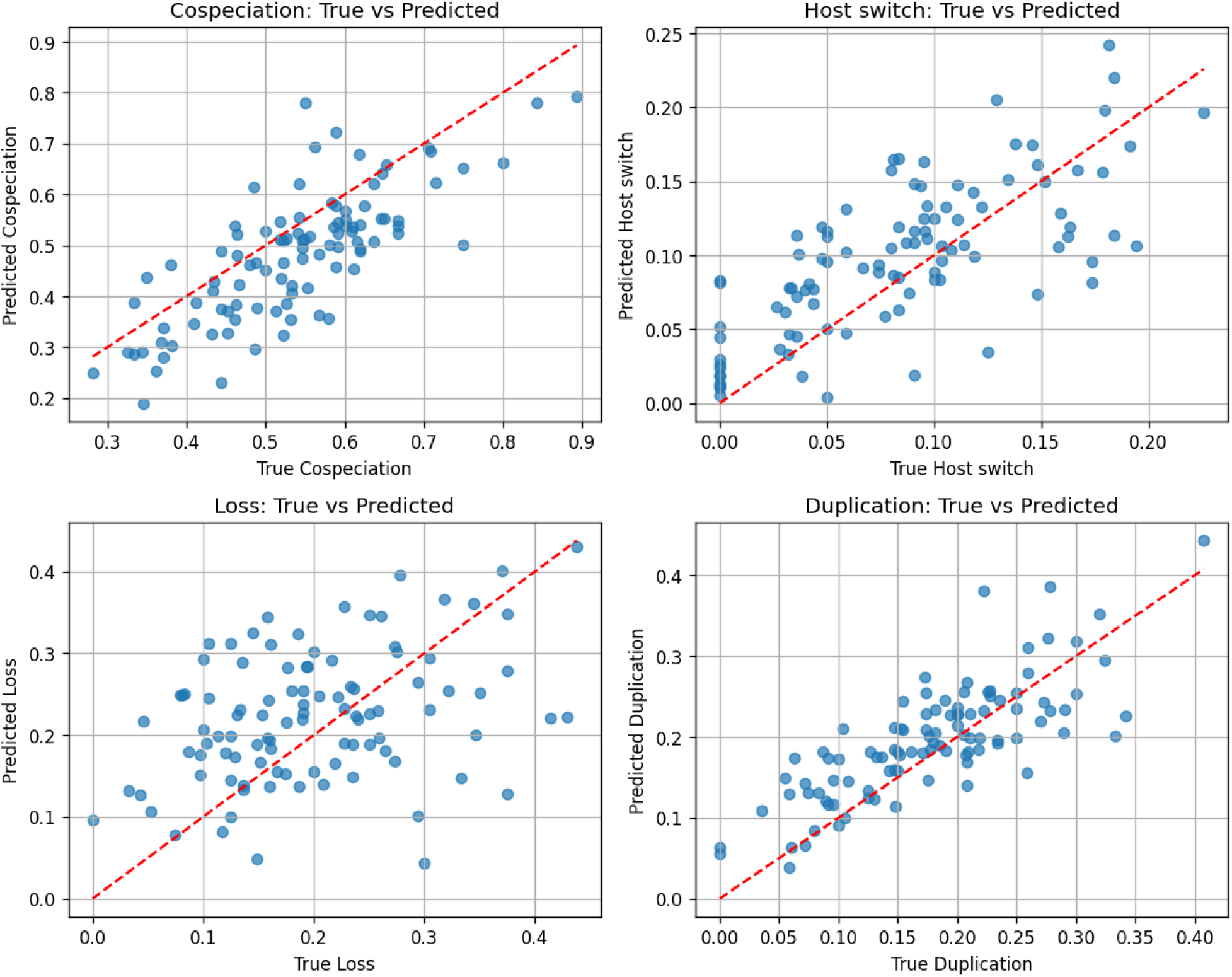
Fox predictions on the treeducken held-out test set (*n* = 100).

### Testing Fox on held-out simulated test data against AmoCoala

We wanted to verify whether Fox was able to accurately recover event frequencies not only on validation split but also on a fully held-out test set coming from the same generator of the training data. Let us recall that the validation split was used to select the model and its parameters while the test set was never used during the training process. Moreover we wanted to test Fox’s ability also on a test set coming from an independent generator not used during the final training of the model (treeducken).

On this second test set we also decided to compare Fox with AmoCoala to see how it compares against other established methods. We chose AmoCoala (Sinaimeri et al. 2023) as the baseline because it targets the same events Fox predicts (the relative frequencies of the four cospeciation, duplication, host-switch and loss events along the symbiont history). AmoCoala estimates the relative frequencies of the four events by ABC: it samples candidate event-probability vectors from a Dirichlet prior, simulates symbiont trees within the observed host tree under each parameter vector, and accepts those whose simulated symbiont trees best match the observed symbiont under a tree-distance statistic, refining the accepted set over successive ABC-Sequential Monte Carlo (SMC) rounds. We ran AmoCoala with *N* = 2000 sampled vectors and *M* = 100 simulated trees per vector, three rounds (*R* = 3) with tolerances *t* = 0.1, 0.25, 0.25, and the default multiple-association metric.

The two models receive very different inputs: AmoCoala requires the host tree, the symbiont tree and the associations map, while Fox does not require trees but MSAs. To make the comparison favorable for the baseline, we supplied AmoCoala with the true simulated host tree, symbiont tree and the association map. Fox instead received the host and symbiont MSAs plus the association map of the two organisms.

The AsymmeTree held-out test set comprises 100 simulated datasets generated with the same pipeline described in paragraph *Tree and sequences simulation* with no overlap with the training or validation data. It spans host trees with 17 to 49 leaves and symbiont trees with 19 to 229 leaves (including lost lineages), with ground-truth frequencies ranging from 0.33 to 0.89 for cospeciation, 0 to 0.24 for host switches, 0 to 0.33 for losses and 0 to 0.37 for duplications. Since this is the simulator Fox was trained on, we use it to measure Fox’s accuracy in its own setting (Table 2 and Fig. 2): Fox recovers event frequencies with MAE 0.016–0.050 and *R*^2^ value 0.45–0.85, and is essentially unbiased (|Bias| *≤* 0.010).

**Table 2.** Fox performance on the AsymmeTree held-out test set.

| | MAE | RMSE | $R^2$ | Pearson $r$ | MRE | Bias |
| --- | --- | --- | --- | --- | --- | --- |
| Cospeciation | 0.0503 | 0.0700 | 0.648 | 0.810 | 0.081 | +0.010 |
| Host-Switch | 0.0160 | 0.0195 | 0.847 | 0.922 | 0.192 | -0.003 |
| Loss | 0.0339 | 0.0455 | 0.449 | 0.688 | 0.305 | -0.005 |
| Duplication | 0.0207 | 0.0283 | 0.827 | 0.910 | 0.112 | -0.002 |

Comparing the two methods on AsymmeTree data would, however, favour Fox. The second test set therefore comprises 100 datasets generated with treeducken (Dismukes and Heath 2021), an independent simulator in which host and symbiont trees evolve jointly. All four events are labelled: host death is set to zero (as in AsymmeTree, whose host trees keep no extinct lineages), symbiont speciation without the host is counted as duplication, symbiont extinction as loss, and an event is counted only if both descendant lineages survive, as in AsymmeTree. Its rates are drawn from the same ranges as Fox’s training data (cospeciation 0.5–1.2, duplication 0.2–0.4, host switch 0.05–0.3, loss 0.2–0.4). This set spans host trees with 15 to 49 leaves and symbiont trees with 17 to 80 leaves (including lost lineages), with ground-truth frequencies ranging from 0.28 to 0.89 for cospeciation, 0 to 0.23 for host switches, 0 to 0.44 for losses and 0 to 0.41 for duplications. On all 100 treeducken datasets (Table 3 and Fig. 3), Fox’s error roughly doubles compared with the one obtained on AsymmeTree test set (mean MAE 0.060 vs. 0.030), but it still tracks the true frequencies of cospeciation, host switches and duplications (Pearson *r* is 0.72–0.78; *R*^2^ value 0.25–0.55), with a moderate underestimation of cospeciation (Bias = *−*0.057). Losses are the hardest event to transfer (*R*^2^ = *−*0.16, *r* = 0.37), which might be explained by the fact that treeducken produces more losses than the in training data, and through a different mechanism.

**Table 3.** Fox performance on the treeducken held-out test set.

| | MAE | RMSE | $R^2$ | Pearson $r$ | MRE | Bias |
| --- | --- | --- | --- | --- | --- | --- |
| Cospeciation | 0.0818 | 0.0993 | 0.246 | 0.775 | 0.152 | -0.057 |
| Host-Switch | 0.0352 | 0.0431 | 0.410 | 0.716 | 0.416 | +0.014 |
| Loss | 0.0805 | 0.1006 | -0.164 | 0.366 | 0.404 | +0.024 |
| Duplication | 0.0424 | 0.0531 | 0.546 | 0.783 | 0.238 | +0.019 |

AmoCoala was favored also in the choice of the subset for the comparison since we selected a smaller subset only based on AmoCoala performances and efficiency. AmoCoala’s inference does not scale easily with tree size: it completed 92 of the 100 treeducken datasets within our compute budget, and the 8 it did not finish are the largest ones (40 to 49 host leaves and 57 to 80 symbiont leaves, against medians of 19 and 29 for the completed datasets). We therefore report the comparison on these 92 datasets, with host trees of 15 to 36 leaves and symbiont trees of 17 to 56 leaves (including lost lineages). AmoCoala’s per-event frequencies were averaged over the 50 accepted solutions of the last round and normalized to relative frequencies to make them directly comparable with Fox’s output. On these 92 datasets (Table 4 and Fig 4), Fox more than halves AmoCoala’s error (mean MAE 0.060 vs. 0.131). Fox is more accurate on cospeciation, host switches and duplications, where AmoCoala’s estimates are essentially uncorrelated with the truth (|*r*| *≤* 0.20) and strongly biased (host switches over-estimated by +0.147, duplications under-estimated by *−*0.142). Losses are the only event on which the two methods are comparable (MAE 0.080 vs. 0.082, Pearson *r* 0.39 vs. 0.36).

**Table 4.** Fox and AmoCoala performance on the 92 treeducken test datasets that AmoCoala completed.

| | MAE | RMSE | $R^2$ | Pearson $r$ | MRE | Bias |
| --- | --- | --- | --- | --- | --- | --- |
| FOX ($n = 92$) | | | | | | |
| Cospeciation | 0.0807 | 0.0980 | 0.297 | 0.778 | 0.150 | -0.054 |
| Host-Switch | 0.0367 | 0.0445 | 0.399 | 0.709 | 0.438 | +0.015 |
| Loss | 0.0799 | 0.1007 | -0.084 | 0.385 | 0.398 | +0.020 |
| Duplication | 0.0433 | 0.0543 | 0.550 | 0.784 | 0.243 | +0.019 |
| AMOCOALA ($n = 92$) | | | | | | |
| Cospeciation | 0.1462 | 0.1820 | -1.421 | -0.195 | 0.272 | +0.049 |
| Host-Switch | 0.1516 | 0.1719 | -7.956 | -0.066 | 1.808 | +0.147 |
| Loss | 0.0823 | 0.1090 | -0.271 | 0.359 | 0.410 | -0.054 |
| Duplication | 0.1430 | 0.1627 | -3.046 | 0.168 | 0.802 | -0.142 |

**Table 5.** Estimated cophylogenetic event frequencies for the cockroach– *Blattabacterium* system (52 host–symbiont pairs). Fox was run on the complete alignments (3,736 host and 35,310 symbiont columns); AmoCoala values are the mean *±*standard deviation over its 50 accepted parameter vectors.

| Event | FOX | AMOCOALA | Difference |
| --- | --- | --- | --- |
| Cospeciation | 0.7048 | $0.5426 \pm 0.251$ | +0.162 |
| Host Switch | 0.1567 | $0.2879 \pm 0.184$ | −0.131 |
| Loss | 0.1056 | $0.1451 \pm 0.079$ | −0.040 |
| Duplication | 0.0329 | $0.0245 \pm 0.024$ | +0.008 |

**Figure 4.**
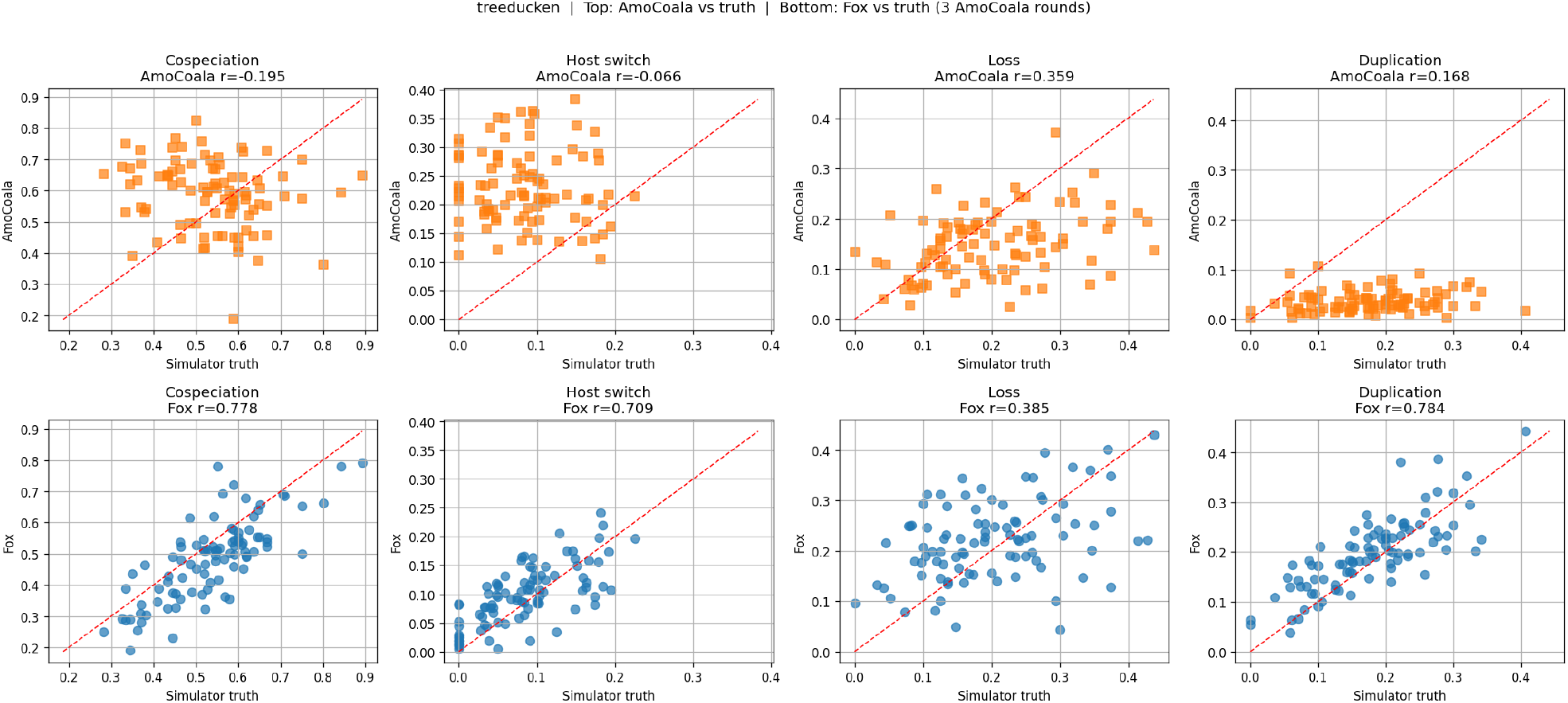
Fox and AmoCoala performance on the 92 treeducken test datasets that AmoCoala completed.

Inference and evaluation for both methods were run on a single MacBook Pro (Apple M4 Pro, 14 cores [10 performance + 4 efficiency], 24 GB unified memory). The runtime gap was not small: Fox completed all 100 predictions in about 13 seconds, whereas AmoCoala required a median of 13 minutes per dataset (up to 74 minutes), about 23.5 hours in total for the 92 datasets it completed, representing a speedup for Fox of more than three orders of magnitude. The difference is structural rather than incidental: AmoCoala must simulate many candidate symbiont trees and compare each to the observed pair to approximate the posterior over event frequencies, whereas Fox produces each prediction in a single forward pass, without the upstream tree-inference step AmoCoala presupposes. We acknowledge that even though Fox is much faster at inference than AmoCoala, its training phase required approximately 300 GPU-hours on 4×H100 GPUs, a cost which an ABC-based method like AmoCoala does not require.

These results suggest that direct inference from MSA pairs can provide more accurate estimates of cophylogenetic event frequencies than a tree-based ABC method, also on data from a simulator it was not trained on, at a much lower inference cost for its user.

### Testing Fox on real data against AmoCoala

Benchmarks on simulated data (Section *Testing* Fox *on held-out simulated test data against* AmoCoala) showed that Fox is able to recover co-phylogenetic event frequencies accurately on data where the ground truth is known. We wanted to assess whether this behavior transfers also to empirical data. Thus, we applied Fox to a real host-symbiont system and compared its output against AmoCoala.

The dataset used is the cockroach–*Blattabacterium* system of Arab et al. (2020), comprising cockroaches, treated as hosts, and their obligate intracellular endosymbiont *Blattabacterium cuenoti*, treated as symbionts. The data consists of 55 hosts sampled across all cockroach families, each cockroach species being linked to the *Blattabacterium* strain isolated from it (55 symbionts in a one-to-one association with their hosts), together with amino-acid alignments of 13 mitochondrial proteins for the hosts and 104 protein-coding genes for the symbionts. Three pairs whose symbiont sequence could not be assigned unambiguously were removed and thus we run both Fox and AmoCoala on the remaining 52 pairs. The concatenated host alignment has 3,736 columns and the concatenated symbiont alignment 35,310 columns. By default Fox reads the complete alignments, whatever their length, even though it was trained on alignments of 250 columns; it then warns that the input lies outside its training range. To show how the estimates depend on the number of columns read, Supplementary Table 6 reports Fox’s estimates and run time on the real dataset, from the first 250 columns up to the complete alignments.

**Table 6.**
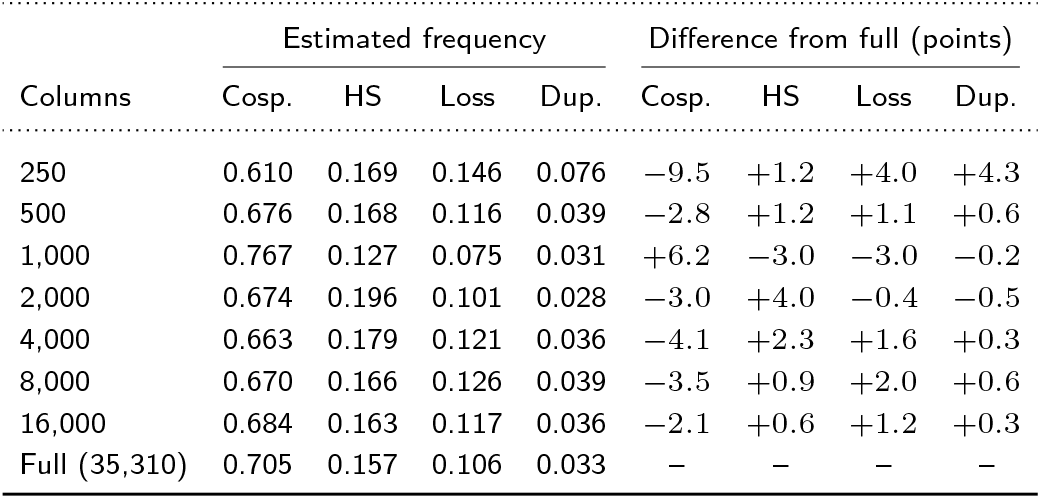
Fox on the cockroach–*Blattabacterium* data (52 pairs) as a function of the number of alignment columns read, and difference from the estimate on the complete alignments. Differences are in percentage points.

| Columns | Estimated frequency |  |  |  | Difference from full (points) |  |  |  |
| --- | --- | --- | --- | --- | --- | --- | --- | --- |
|  | Cosp. | HS | Loss | Dup. | Cosp. | HS | Loss | Dup. |
| 250 | 0.610 | 0.169 | 0.146 | 0.076 | −9.5 | +1.2 | +4.0 | +4.3 |
| 500 | 0.676 | 0.168 | 0.116 | 0.039 | −2.8 | +1.2 | +1.1 | +0.6 |
| 1,000 | 0.767 | 0.127 | 0.075 | 0.031 | +6.2 | −3.0 | −3.0 | −0.2 |
| 2,000 | 0.674 | 0.196 | 0.101 | 0.028 | −3.0 | +4.0 | −0.4 | −0.5 |
| 4,000 | 0.663 | 0.179 | 0.121 | 0.036 | −4.1 | +2.3 | +1.6 | +0.3 |
| 8,000 | 0.670 | 0.166 | 0.126 | 0.039 | −3.5 | +0.9 | +2.0 | +0.6 |
| 16,000 | 0.684 | 0.163 | 0.117 | 0.036 | −2.1 | +0.6 | +1.2 | +0.3 |
| Full (35,310) | 0.705 | 0.157 | 0.106 | 0.033 | – | – | – | – |

Fox was run directly on the two MSAs and their mappings. AmoCoala requires the host and symbiont trees, which Arab et al. (2020) provide only as a figure: we transcribed the topologies of their maximum-likelihood trees (Figure 1 of Arab et al. 2020) for the same 52 host–symbiont pairs as Fox with the same one-to-one associations; it was run under the following settings: `-N 2000 -M 100 -R 3 -t 0.1,0.25,0.25`.

Table 5 and Figure 5 report the two sets of estimates. Both methods identify cospeciation as the most frequent event, consistent with the vertical transmission of *Blattabacterium*, and both rank the four events in the same (decreasing) order of importance: cospeciation, host switching, loss and then duplication. Run on the complete alignments, Fox assigns 70.5% to cospeciation, 15.7% to host switching, 10.6% to loss and 3.3% to duplication. AmoCoala puts less weight on cospeciation (54.3%) and more on host switching (28.8%), with 14.5% to loss and 2.5% to duplication. Across its 50 accepted parameter vectors, AmoCoala’s estimates vary widely, with standard deviations of 0.25 for cospeciation, 0.18 for host switching, 0.08 for loss and 0.02 for duplication; all four Fox estimates fall within one standard deviation of AmoCoala’s mean. Reading the complete alignments (3,736 host and 35,310 symbiont columns, far beyond the 250 columns used in training) took Fox about 6 minutes and 16 GB of memory on CPU, while AmoCoala took about 20 minutes to complete the 3 rounds. With 250 columns, Fox returns its estimate in under 1 second. Compared with the full-alignment estimate, (Table 6) this approximation assigns 9.5% of the events to a different category (cospeciation *−*9.5, host switching +1.2, loss +4.0 and duplication +4.3 percentage points), and preserves the ranking of the four events. Reading 2,000 columns, which takes 5 seconds, lowers this difference to 4% (Supplementary Table 6).

**Figure 5.**
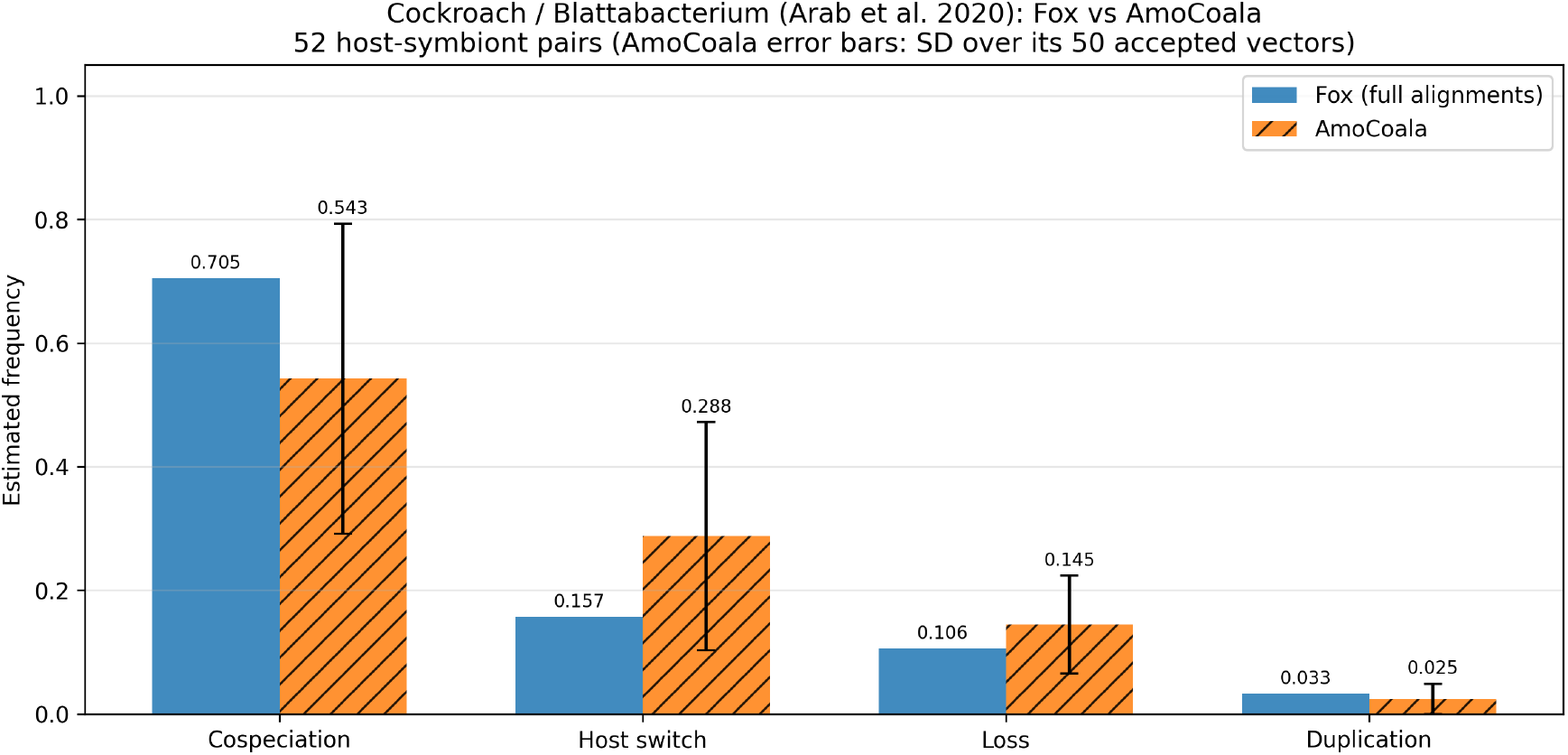
Fox versus AmoCoala event-frequency estimates on the cockroach–*Blattabacterium* dataset. Fox was run on the complete alignments (3,736 host and 35,310 symbiont columns); AmoCoala error bars show the standard deviation over its 50 accepted vectors.

### Sensitivity to MSA length and host-symbiont associations

We first wanted to assess sensitivity of Fox with respect to MSA lengths. Indeed, Fox was trained with MSAs comprising 250 columns, but can be used with any MSA length. A sequence length analysis (Supplementary Figure 5 and Supplementary Table 3) shows that Fox’s accuracy is essentially stable when the MSAs are shortened from 250 to 150 columns, degrades at 100 columns, and falls off sharply at 50 columns, where cospeciation, host switches and losses become unreliable. Nonetheless duplication is the most robust event, keeping high *R*^2^ even at 50 columns.

Finally, we tested whether Fox uses the host–symbiont associations. On the 100 held-out AsymmeTree test datasets, we shuffled the links between hosts and symbionts while keeping the sequences unchanged (Supplementary Table 2). The predicted cospeciation frequency drops in every test dataset, from 0.63 to 0.23 on average, and on the cockroach data from 0.61 to 0.39, while host switching and loss increase. Fox’s predictions therefore depend on which symbiont is associated with which host, and not only on the two alignments.

## Methods

### Supervised learning in simulation-based inference

We rely on simulation-based inference (Cranmer et al. 2020): a family of methods that is used to estimate parameters *θ* of a probabilistic generative model *p*(*x* | *θ*) when the likelihood is unavailable but the model can be used to produce samples *x ~ p*(*·* | *θ*). In our scenario, *θ* is the vector of four relative event frequencies (cospeciation, host-switch, loss, duplication) that governs a coevolution process while *x* is the pair of host and symbiont MSAs that results from running that process and then evolving sequences along the resulting trees, together with their leaf mapping. Our goal is not to evaluate or find *p*(*x* | *θ*) but instead, we generate many pairs (*θ*_*i*_, *x*_*i*_) by simulation and use them as labeled training data, generating inference of the *θ* from *x* as a supervised regression problem. We simulate data (MSAs and host-symbiont associations) under a probabilistic coevolution model using various rates covering the space of regimes the model is later expected to generalize across. Similarly to ABC, our method is likelihood-free, in that it relies on a probabilistic model framework whose likelihood is never computed. However a major difference with ABC lies in that the latter needs simulations that are specific to each real dataset at hand, resulting in computationally intensive tools (though less intensive than Monte-Carlo Markov Chains likelihood computation, for e.g.). On the contrary, our approach learns the function that links the data with the true parameters and requires simulations only prior to a learning phase. Then once the model has been trained on these simulated data, the user can employ the tool on any new dataset without the need to produce new synthetic data and thus at almost no cost. This is what is usually referred to as amortized inference where the computational cost is paid only once during training and then every subsequent dataset is processed in a single forward pass at inference time. ABC methods instead pay a comparable simulation cost per input dataset since each new host-symbiont pair requires drawing fresh candidate trees under the prior and comparing them to the observation. Our model thus trades a large one time training cost for a near zero marginal cost per prediction whereas ABC trades no training cost for a marginal cost that scales with every new dataset analyzed.

We also want to emphasize what each method uses and what it does not need to know. Fox is only given the host and symbiont MSAs, the host-symbiont sequence mapping, and the pairwise evolutionary distances derived from those alignments; it is never given the true host or symbiont tree topology, branch lengths, or the underlying event rates used to simulate the data; these rates or rather their normalized relative frequencies, are precisely the quantities that the model is trained to recover. This stands in contrast to tree-based ABC approaches, which require the host and symbiont trees (or a tree-reconstruction step) as input alongside the association map.

For the generation of co-evolved host and symbiont trees we used AsymmeTree (Schaller et al. 2022) for the training data while treeducken (Dismukes and Heath 2021) was used only to generate an independent test set. We combined these with a sequence evolution model (LG+Γ+invariable-sites) along each tree, to generate host and symbiont amino acids MSAs. With these data at hand, we have designed an elaborate neural network architecture that learns the frequencies of the four co-evolutionary events.

### Network architecture

Fox (Fig. 6) is a transformer-based model for predicting cophylogenetic event frequencies from paired host and symbiont multiple sequence alignments. It follows the multi-head attention framework (Vaswani et al. 2017), takes MSAs as the primary phylogenetic input (similarly to Phyloformer, Nesterenko et al. 2025), and incorporates the idea of jointly refining sequence-level representations and representations between pairs of sequences. Indeed, the latter already proved useful to infer phylogenetic trees from sequences relying on neural networks (Blassel et al. 2026). The model receives a host MSA, a symbiont MSA and the known host-symbiont sequence associations. From the MSAs, it will also compute host and symbiont pairwise distance matrices. The output is a normalized four-dimensional vector for cospeciation, host-switch, loss, and duplication frequencies obtained with a softmax function so it is directly comparable to normalized event-frequency labels.

**Figure 6.**
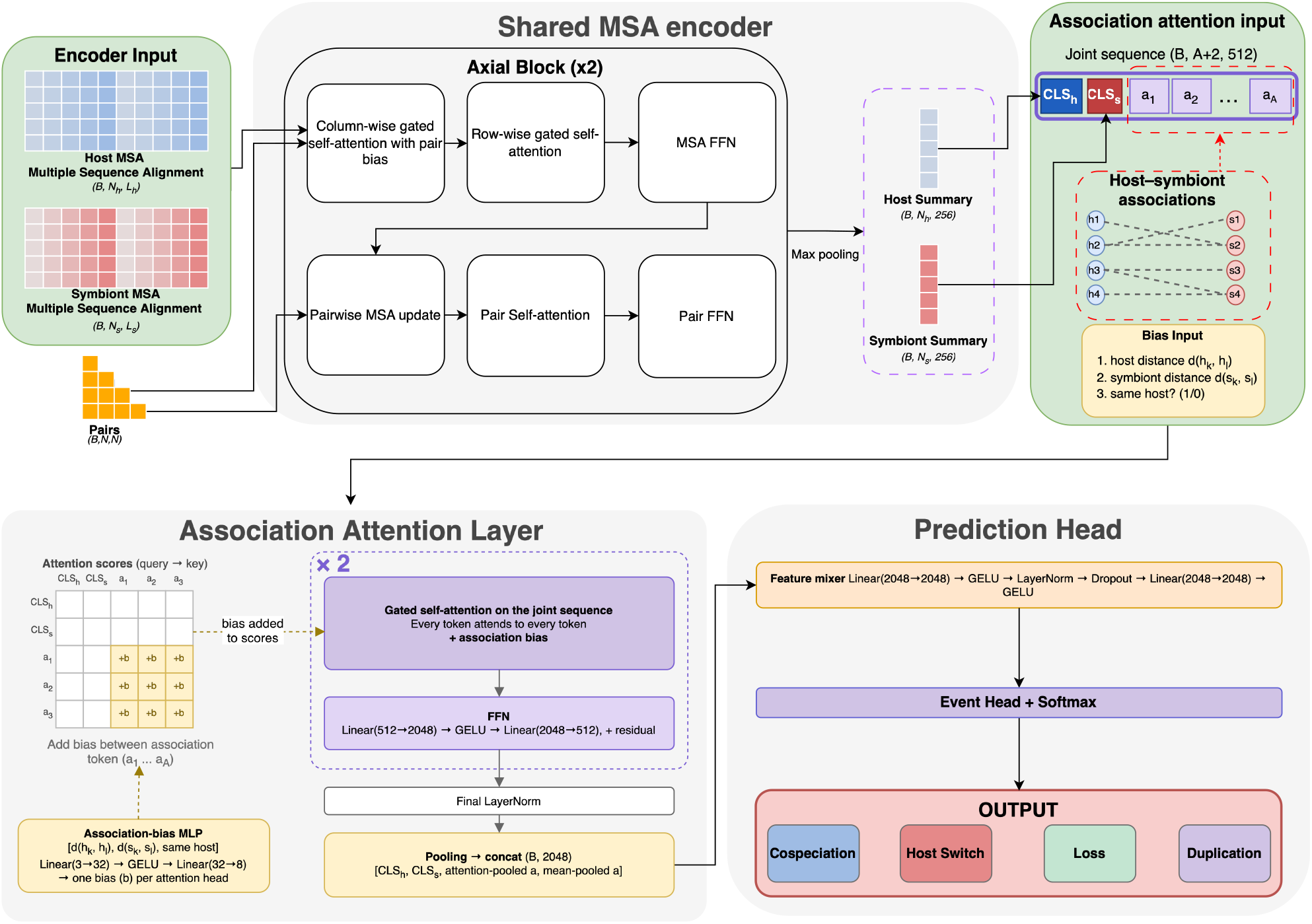
Architecture of the Fox model for host–symbiont coevolutionary event prediction. Inputs: host (B, *N*_*h*_, *L*_*h*_) and symbiont (B, *N*_*s*_, *L*_*s*_) MSAs, a within-group pair tensor for each (host (B, *N*_*h*_, *N*_*h*_), symbiont (B, *N*_*s*_, *N*_*s*_)) computed from the respective MSA, and the host–symbiont mapping. A shared MSA encoder processes each alignment through axial blocks combining column-wise gated self-attention with pair bias, row-wise gated self-attention and an MSA FFN, alongside a pair track; in each block the pair track biases the MSA column attention and the MSA track updates the pair track. The encoder outputs per-sequence summaries (B, N, 256) and a global classification token (CLS; B, 512) for host and symbiont. For each of the A (host–symbiont associations) in the mapping, an association token (B, A, 512) is built from the summaries of its host and symbiont sequences. Association attention layers mix the two CLS tokens with the A association tokens; attention between two associations is biased by their host distance, symbiont distance and whether they share a host. The two CLS tokens and the attention- and mean-pooled association tokens are concatenated (B, 2048) and passed through a feature mixer (2× Linear–GELU with LayerNorm/Dropout). An event head with softmax outputs probabilities for the four events: cospeciation, host-switch, loss and duplication. Shapes: B batch size; *N*_*h*_, *N*_*s*_ number of host/symbiont sequences; *L*_*h*_, *L*_*s*_ alignment lengths; A number of host–symbiont associations

In what follows, “within-groups” pairs of sequences refer to either pairs of host or pairs of symbiont sequences. It is to be distinguished from host-symbiont associated sequences. The architecture has three main stages:

- **a shared MSA encoder** (Blassel et al. 2026) applied separately to the host and symbiont alignments, built from two axial blocks that alternate gated (Qiu et al. 2025) within-group pair-based column attention, row attention (on each individual sequence), and self-attention on a within-group pair-level representation (not focusing on columns separately);
- two-layer **association-aware attention** over the host and symbiont summaries and one token per host–symbiont association, with attention between associations biased by host and symbiont distances;
- a four-event rate **prediction head** conditioned on the encoding produced by the previous stages.

The reference instantiation uses hidden dimension *d* = 256 for encoding each MSA, within-group pair encoding dimension 64, association token dimension *d*_cls_ = 512, together with 8 attention heads, 2 axial encoder layers and 2 association attention layers.

### Shared MSA encoder

Fox consists of a single `MSAEncoder` (Fig. 6 Shared MSA encoder) that is called once on the host MSA and once on the symbiont MSA, so both alignments are encoded with shared weights rather than two independent encoders. Host-specific and symbiont-specific information is introduced only after encoding through separate host/symbiont sequence summaries, global classification token descriptors (CLS), and token-type embeddings Each MSA is represented as a tensor (*B, N, L*), where *B* is the batch size, *N* is the number of sequences, and *L* is the alignment length (with *N* = *N*_*h*_, *L* = *L*_*h*_ for the host and *N* = *N*_*s*_, *L* = *L*_*s*_ for the symbiont). Tokens use a vocabulary of size 23, comprising 20 amino acids, a gap token, an unknown token, and a padding token. The token IDs are one-hot encoded and passed through a bias-free linear projection followed by ReLU (Rectified Linear Unit), producing per-site embeddings in R*d* with *d* = 256. In parallel, the encoder maintains a within group pair track of shape (*B, N, N*, 64) (with *N* equal to *N*_*h*_ or *N*_*s*_) over all sequence pairs within each group (in Fig. 6 the orange “Pairs”). Pair features are initialized by projecting the token representation, averaging along sequence positions while ignoring padding, and combining sequence summaries with an outer-sum operation. Each scalar distance is also projected through a multi-layer perceptron (MLP) and added to the pair representation. We pre-compute these distances from the tokenized MSAs using a Jukes-Cantor estimated evolutionary distance (Jukes and Cantor 1969) with padding-aware pairwise comparisons; for each pair we count mismatches over jointly non-padded columns, and set 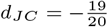 ln 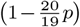, the Jukes–Cantor distance for 20 amino-acid states, with *p* being the proportion of differing residues for that pair of sequences. We clamp *p* to [0, 0.94] to keep the logarithm argument positive, since *d*_*JC*_ diverges as *p →* 0.95 (the protein Jukes–Cantor saturation point) and is undefined beyond it.

### Axial encoder block

The embeddings and the pair track are refined by a stack of two axial blocks (Fig. 6, *Axial Block (×2)*), each an EvoPF-style (Blassel et al. 2026) unit, the latter being a variation of EvoFormer (Jumper et al. 2021) that update the two tracks in sequence.

The MSA track is updated in three steps: (i) *column-wise gated self-attention with pair bias*: gated column attention across sequences at each alignment position (site or residue); the attention logits receive a learned per-head bias from the current pair representation; (ii) *row-wise gated self-attention*: gated attention along residue positions within each sequence; (iii) a position-wise *MSA feed-forward* update.

The within-group-pair track is updated from the already updated MSA: the *pairwise MSA update* mean-pools the normalized MSA features over sequence positions, projected into two intermediate representations, and combined through summed and absolute-difference within-group-pair features. An outer-product-mean update is available as an alternative, but it is disabled in the final model to make for efficiency reasons. The update is followed by within-group *pair self-attention* and a within-group *pair feed-forward* network (FFN), both with residual dropout.

The two tracks (MSA and within-group-pair) thus exchange information in a single directed way per block: the pair track influences the MSA track only through the per-head bias in step (i) (the *with pair bias* arrow), while the MSA track feeds back into the pair track through the pairwise MSA update. In the final block the pair update is disabled, since no subsequent column-attention layer consumes it.

### Sequence summaries

After the last encoder block, the residue dimension is reduced with masked max-pooling, to one contextualized embedding per sequence (*B, N*, 256). A learned query token then attends over all sequences embeddings (multi-head attention pooling, ignoring padded sequences), producing a global MSA summary. The encoder returns both the per-sequence embeddings, used for host-symbiont mapped-pair fusion, and the global summary vector (CLS), used as the first tokens of the association attention sequence (Fig. 6, top right corner).

#### Association-aware attention

Fox uses association-aware attention, in which each host–symbiont link becomes one token, described below. For each batch element, the known host-symbiont associations are represented as index pairs. The model gathers the corresponding host and symbiont sequence embeddings from the shared MSA encoder outputs. These embeddings are concatenated and passed through a two-layer MLP to a single association token of dimension *d*_cls_ = 512, so each token stands for one host–symbiont link. The model constructs a sequence where the first two tokens are the host and the symbiont global MSA summaries (CLS) and the remaining tokens are the association tokens, each with a learned token-type embedding (host CLS, symbiont CLS, association) (Fig. 6). A mask prevents padded association tokens (added so that systems with different numbers of associations fit in one batch) from contributing to attention and pooling.

Two layers (Fig. 6, *Association Attention Layer*) fuse the host and symbiont information. At each layer, every token attends to all tokens of the sequence, so host summaries, symbiont summaries and association tokens exchange information in both directions. The attention between two association tokens receives a learned per-head bias, computed by a small MLP from the distance between their host sequences, the distance between their symbiont sequences (both taken from the Jukes–Cantor matrices) and whether they share the same host, so that the model can directly compare whether symbiont distances follow host distances. Each layer has its own attention module, layer normalizations, and feed-forward network.

The model then builds a fixed-size joint descriptor. It keeps the updated host CLS and symbiont CLS tokens, and pools the association tokens in two ways: a masked softmax pooling with a learned scalar scoring function, and a masked mean. These four vectors are concatenated, giving a representation of size 4*d*_cls_ = 2048.

#### Prediction head

The concatenated representation is processed by a feature-mixing MLP with GELU (Gaussian Error Linear Unit) activations, layer normalization, and dropout (Fig. 6, Bottom right).

The head emits four real scores *z* = (*z*_cosp_, *z*_hs_, *z*_loss_, *z*_dup_); a softmax maps them to

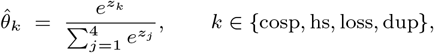

forcing the four outputs to be positive and to sum to one. The prediction therefore lies on the same simplex as the target *θ* and is directly comparable to it, with no post-processing.

#### Tree and sequences simulation

There exist multiple models able to generate synthetic co-phylogenetic data (Di Palma et al. 2026); our only constraint is that we need dated trees since branch lengths are fundamental for the generation of MSAs. For this reason, we had to exclude some models as possible solution for generating our training data. We decided to train our model on AsymmeTree (Schaller et al. 2022) and then test the performance on data generated by another generator such as treeducken (Dismukes and Heath 2021). Note that AsymmeTree is initially designed in the gene-species context but it can be used in the host-symbiont framework as explained in Di Palma et al. (2026). The two generators differ in the way host and symbiont histories are produced. AsymmeTree first samples a host tree relying on a birth-death process conditioned on the number of leaves and then simulates symbiont evolution with explicit cospeciation, host-switch (called Horizontal Gene Transfer, HGT), duplication, and loss rates. In our AsymmeTree runs, host leaf counts were sampled in the range *N*_*h*_ *∈* [15, 50], host birth and death rates were drawn from broad uniform priors and we used one of five evolutionary ages *τ ∈ {*1.5, 2, 2.5, 3, 3.5*}* (see details in Table 7). Symbiont event rates were sampled independently. For each replicate we counted the realized co-evolutionary events along the symbiont history and normalized them by the total event count, yielding a target vector on the simplex (entries non-negative, summing to one). Predicting relative frequencies rather than raw counts decouples the target from tree size and makes replicates of different scales directly comparable. By contrast, treeducken co-simulates host and symbiont histories under a time-conditioned birth-death cophylogenetic process. Host and symbiont leaf counts are therefore emergent rather than fixed in advance. The simulator directly models a cospeciation rate, independent host and symbiont birth rates, coupled death rates, and host-expansion or host-switch events. treeducken records six event types along the symbiont tree, which do not map one-to-one onto the four events of AsymmeTree: in particular, with non-zero host death, symbionts are also lost when their host goes extinct. For the independent test set (Section *Testing* Fox *on held-out simulated test data against* AmoCoala) we built an explicit mapping to all four events: host death is set to zero, symbiont speciation without the host is counted as duplication, and each extinct symbiont lineage as one loss, following the same counting convention as AsymmeTree.

For both simulators (training and test data), amino-acid MSAs of length *L* = 250 were generated with AliSim (Ly-Trong et al. 2022) under the same LG+Γ+invariable-sites setting. Each model received paired host and symbiont MSAs, the host-symbiont sequence mapping, and Jukes-Cantor distance matrices computed on the MSAs for both the host and the symbiont.

We generated roughly 1.5 million pairs of trees using the different input parameters found in Table 7 with a standard 80/20 training-validation split for training the model. The trees we keep, and hence the MSAs and the association map, contain only what real data can show: symbiont lineages lost before the present are pruned and the planted root edge is removed.

**Table 7.** Parameter priors used to simulate the training data with AsymmeTree. A cospeciation event occurs at each host speciation, while host switch, duplication and loss are explicit symbiont event rates. Each replicate draws its parameters independently from these priors.

| Parameter | Prior |
| --- | --- |
| Host leaves $N_h$ | $\mathcal{U}[15, 50]$ |
| Host birth $\lambda_h$ | $\mathcal{U}[0.5, 1.2]$ |
| Host death $\mu_h$ | $\mathcal{U}[0.2, 0.4] \cdot \lambda_h$ |
| Host switch $r_{hs}$ | $\mathcal{U}[0.05, 0.30]$ |
| Duplication $r_{dup}$ | $\mathcal{U}[0.20, 0.40]$ |
| Loss $r_{loss}$ | $\mathcal{U}[0.20, 0.40]$ |
| Age $\tau$ | $\{1.5, 2, 2.5, 3, 3.5\}$ |

Empirical distributions of the event frequencies in the training data are shown in Supplementary Figure 1, and the corresponding tree-structure distributions in Supplementary Figure 2.

#### Training Setup

Fox was trained on the Jean Zay HPC cluster (IDRIS/CNRS) using 4 *×* NVIDIA H100 80 GB GPUs on a single node (96 CPU cores in total), via Distributed Data Parallel (DDP) with PyTorch 2.6.0 and Lightning Fabric. Total training took approximately 300 GPU-hours across these 4 GPUs. All runs used a fixed random seed (42) for reproducibility.

The model follows the reference instantiation of Sec. *Network architecture* (approximately 21.5 million trainable parameters; full configuration in Supplementary Table 1). Column attention used a fused pair-biased FlexAttention kernel, and the pair track was initialized from both the MSA content and the Jukes–Cantor distance matrix.

Training was allowed up to 35 epochs (early stopping ended it after epoch 26, and we release the best validation checkpoint (epoch 23)), a per-GPU batch size of 32 and gradient accumulation over 4 steps, giving an effective batch size of 512. Parameters were optimized with AdamW (lr = 10^*−*4^, *β*_1_ = 0.9, *β*_2_ = 0.999, weight decay = 0.05). The learning rate followed a cosine schedule with linear warm-up over the first 15 % of training steps, decaying to 10 % of the peak rate. The objective was a weighted squared error applied independently to each event-frequency output:

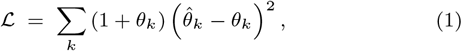

where *k ∈ {*cosp, hs, loss, dup*}*, 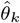 is the predicted frequency and *θ*_*k*_ the target (Section *Prediction head*). ℒ is computed for each dataset (one host tree, one symbiont tree and their tip associations) and averaged over the datasets in a batch. The weight (1 + *θ*_*k*_) gives in which an event is frequent up to twice the weight of systems in which it is absent, so the model does not learn to underpredict events that are rare in most training systems.

A shape-bucketed sampler grouped sequences by (host taxa, symbiont taxa) with bucket granularity 4 to minimize padding overhead. Validation was run both mid-epoch and at epoch end, and training was stopped early if validation loss did not improve for 3 consecutive epochs (minimum Δ = 0), with the patience counter held during warm-up.

Fox was trained on datasets with at most 50 host leaves, at most 142 symbiont leaves and alignments of at most 250 columns (Table 7). These bounds were chosen when designing the training data, to avoid very small or very large trees and to keep the GPU memory needed per batch within the capacity of the GPUs; they are not limits of the architecture. No layer of the network depends on the number of hosts, the number of symbionts or the alignment length: Fox takes the complete alignments as input, whatever their size, and shorter alignments are simply padded. When an input exceeds any of the training bounds, Fox runs on it in full and issues a warning that the input lies outside its training range, since its accuracy there has not been evaluated systematically. Time and memory grow with the size of the input: on the cockroach–*Blattabacterium* data (Section *Testing* Fox *on real data against* AmoCoala), reading the complete alignments (3,736 host and 35,310 symbiont columns) took about 6 minutes and 16 GB of memory on CPU, against under 1 second for 250 columns. The effect of alignment length on accuracy is analysed in Supplementary Figure 5.

## Conclusion

We introduced Fox, a transformer that infers the relative frequencies of the four cophylogenetic events (cospeciation, host-switch, loss and duplication) directly from a pair of host and symbiont multiple sequence alignments and their host-symbiont mapping without any tree reconstruction. Fox pays its computational cost once, during training, and then processes any new host-symbiont system in a single forward pass. To assess its accuracy and generalizability, we evaluated Fox on three types of data: held-out synthetic data from the training simulator (MAE 0.016–0.050, *R*^2^ 0.45–0.85), synthetic data from an independent simulator (roughly doubled errors, but 45–76% lower MAE than AmoCoala for cospeciation, host switches, and duplications, with similar accuracy for losses), and one empirical cockroach–*Blattabacterium* dataset (recovering a cospeciation-dominated profile consistent with AmoCoala), indicating that patterns learned from simulated data also transfer to real sequences. Consistent with its single-pass design, Fox was over three orders of magnitude faster than AmoCoala, completing datasets that the latter could not handle.

Fox still has some limitations. It is trained under a specific sequence-evolution model (LG+Γ+invariable-sites) and on a limited range of tree sizes and event regimes, so its performance outside these settings remains to be tested. In particular, future work should assess how robust Fox is when real data deviate from the sequence-evolution model used for training. Losses remain the most difficult event to infer from sequence data alone.

Several extensions follow naturally. First, Fox currently provides only point estimates of event frequencies. Adding calibrated uncertainty estimates would make it possible to distinguish confident predictions from uncertain ones.Second, Fox was trained on alignments of 250 columns, at most 50 hosts and at most 142 symbionts; it runs on larger inputs, but its accuracy there has not been evaluated systematically. Training on longer alignments and larger trees would make Fox reliable on larger and more heterogeneous datasets. Third, Fox currently uses protein alignments only. Extending the model to nucleotide sequences would make it applicable to DNA data. Finally, a longer-term goal is to move beyond estimating global event frequencies and predict *where* individual events occur on the host and symbiont trees.

## Data availability

The Fox source code and the pretrained model are freely available on GitHub at https://github.com/Gabbo240900/Fox. The training data is not distributed given the large numbers of datasets used for training, but we provide the full simulation pipeline used to generate it, so that the datasets can be generated and the model can be retrained from scratch. The simulated datasets and the real cockroach–*Blattabacterium* dataset used for held-out analysis, are instead included in the repository.

## Author contributions statement

BS, CM and LJ conceived the method. GDP and LJ designed and tuned the neural network architecture. GDP conducted the experiments. All authors analyzed the results and wrote and reviewed the manuscript.

## Acknowledgments

This work was granted access to the HPC resources of IDRIS under the allocation 2025-AD010716380 made by GENCI.

Generative AI tools were used for English language editing and code assistance. All scientific content, analyses, interpretations, and conclusions were done by the authors, who take full responsibility for the final manuscript and the code used in this study.

