## Supplementary material for "Fox: Predicting rates of host-symbiont co-evolution events with association-aware attention"

### Training configuration

Supplementary Table 1: Training configuration for the released Fox model

| Setting | Value |
|---|---|
| Hardware | 4× NVIDIA H100 80 GB, single node (Jean Zay, IDRIS/CNRS) |
| CPU cores | 96 |
| Distributed backend | Distributed Data Parallel (DDP), Lightning Fabric |
| Framework | PyTorch 2.6.0 |
| Total compute | ∼300 GPU-hours |
| Trainable parameters | ∼21.5M |
| Axial encoder layers | 2 |
| Association attention layers | 2 |
| Attention heads | 8 |
| Hidden dim $d$ | 256 |
| Within-group pair dim | 64 |
| Association token dim $d_{\text{cls}}$ | 512 |
| Dropout | 0.10 |
| Optimizer | AdamW ($\beta_1$=0.9, $\beta_2$=0.999, weight decay 0.05) |
| Peak learning rate | $10^{-4}$ |
| LR schedule | cosine, 15% linear warm-up, decay to 10% of peak |
| Loss | Huber ($\delta = 1.0$), per event-frequency output |
| Epochs | 35 (early stop, patience 3, min $\Delta = 0$) |
| Per-GPU batch size | 32 |
| Gradient accumulation | 4 steps (effective batch 512) |
| Sampler | shape-bucketed by (host taxa, symbiont taxa), granularity 4 |
| Sequence length $L$ | 250 |
| Random seed | 42 |

### Training and test supplementary analysis

To check that Fox uses the host–symbiont associations, we shuffled the links while keeping the sequences unchanged: the host side of every link was relabelled with a random permutation of the hosts, so each symbiont is attached to a random host while every host keeps its number of symbionts. We ran 20 shuffles on the cockroach–*Blattabacterium* dataset (all 52 pairs) and 20 shuffles on each of the 100 AsymmeTree held-out test datasets. To keep the 21 runs on the cockroach data fast, this test reads the first 250 columns of each alignment, the training length, so its original-links prediction differs from the full-alignment estimate of the main text.

If the model ignored the associations, the predictions would not change. Instead, the predicted cospeciation frequency drops from 0.61 to 0.41 on the cockroach data, and from 0.63 to 0.23 on average on the AsymmeTree test set, where it decreases in all 100 datasets. The freed frequency moves mostly to host switching and loss, which increase in all 100 test datasets (Supplementary Table 2). This is the expected behaviour: once the links no longer follow the phylogenies, the data look less like cospeciation and more like host switches and losses.

Supplementary Table 2: **Predictions of Fox with original and shuffled host–symbiont links.**

| | Cospeciation | Host-Switch | Loss | Duplication |
|---|---|---|---|---|
| *Cockroach–Blattabacterium* | | | | |
| Original links (first 250 columns) | 0.610 | 0.169 | 0.146 | 0.076 |
| Links shuffled (mean ± SD, 20) | $0.406 \pm 0.045$ | $0.223 \pm 0.015$ | $0.205 \pm 0.021$ | $0.166 \pm 0.028$ |
| AsymmeTree *test set, mean over 100 datasets* | | | | |
| Original links | 0.630 | 0.080 | 0.106 | 0.183 |
| Links shuffled (20 per dataset) | 0.230 | 0.263 | 0.303 | 0.205 |
| Datasets where the value decreased | 100/100 | 0/100 | 0/100 | 27/100 |

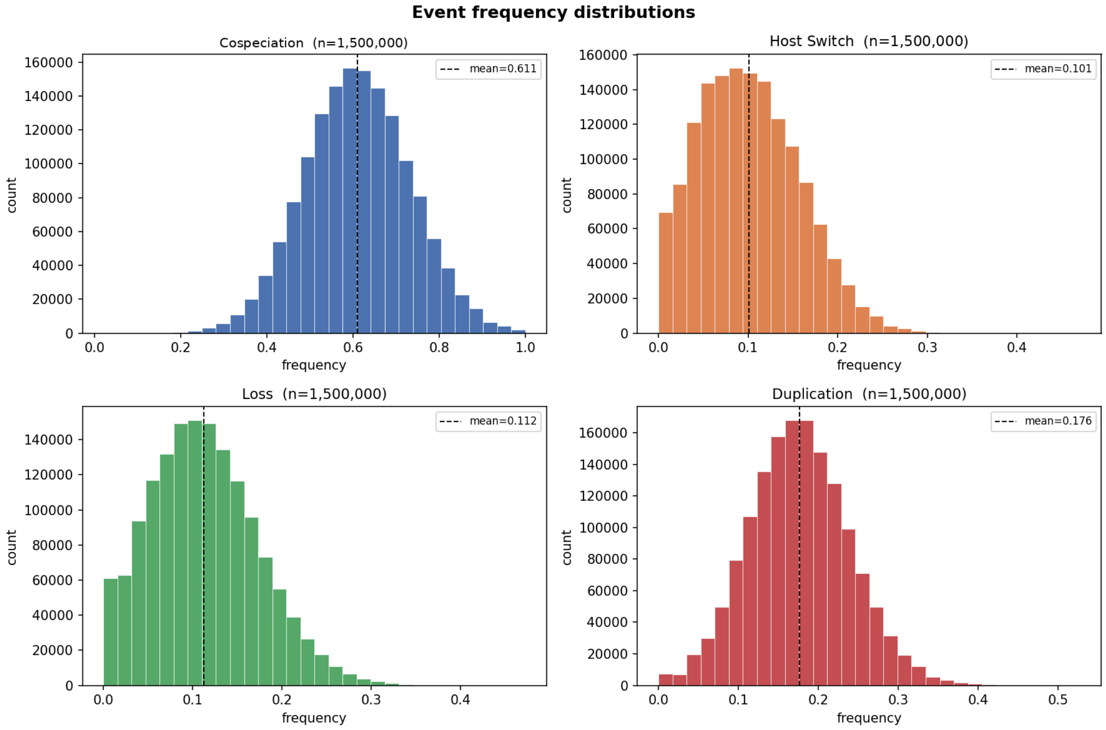


Supplementary Figure 1: Training-data event-frequency distributions.

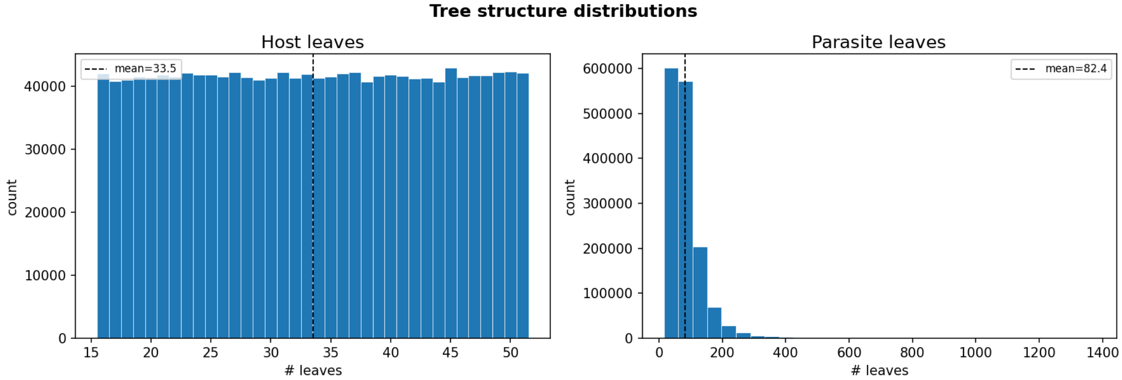


Supplementary Figure 2: Training-data tree-structure distributions.

Supplementary Table 3: Fox accuracy vs. alignment length on the 100 held-out AsymmeTree test datasets

| | MAE | | | | $R^2$ | | | |
|---|---|---|---|---|---|---|---|---|
| Length | Cosp | HS | Loss | Dup | Cosp | HS | Loss | Dup |
| 50 | 0.1280 | 0.0650 | 0.0769 | 0.0276 | −0.546 | −1.114 | −1.211 | 0.728 |
| 100 | 0.0639 | 0.0255 | 0.0419 | 0.0222 | 0.525 | 0.618 | 0.255 | 0.816 |
| 150 | 0.0517 | 0.0189 | 0.0341 | 0.0214 | 0.645 | 0.788 | 0.433 | 0.818 |
| 200 | 0.0515 | 0.0167 | 0.0338 | 0.0210 | 0.651 | 0.835 | 0.439 | 0.829 |
| 250 | 0.0503 | 0.0160 | 0.0339 | 0.0207 | 0.648 | 0.847 | 0.449 | 0.827 |

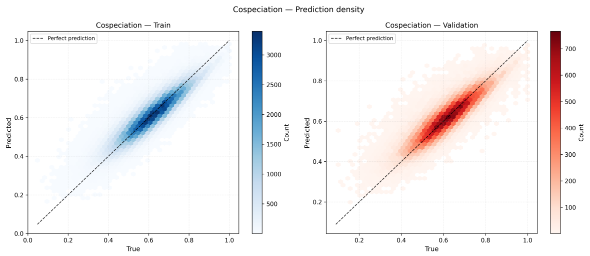


(a) Cospeciation

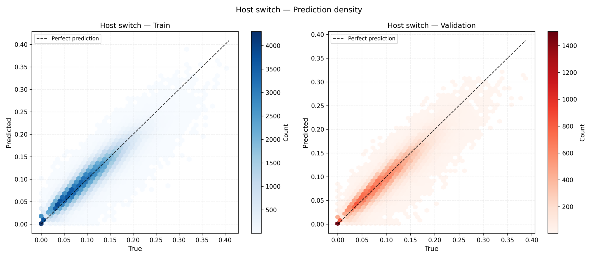


(b) Host Switch

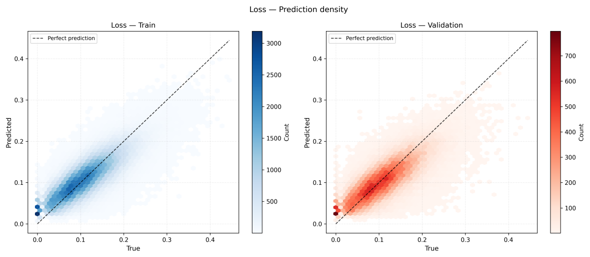


(c) Loss

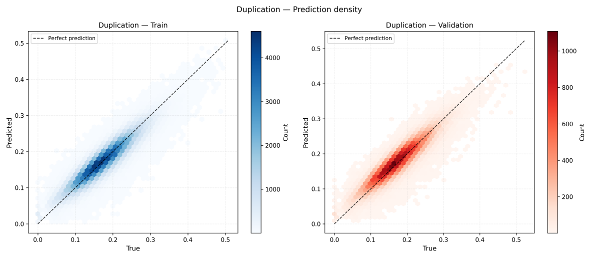


(d) Duplication

Supplementary Figure 3: Two-dimensional density of predicted vs. true frequencies on the training set (left, blue) and validation set (right, red) for the four events.

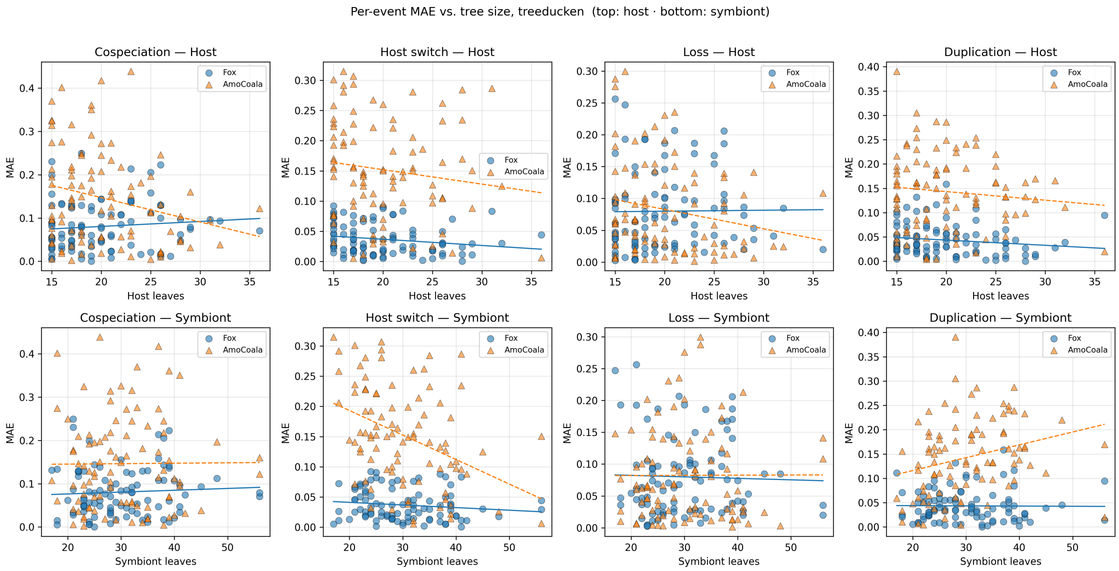


Supplementary Figure 4: AmoCoala vs Fox error as a function of tree size, per event (scatter, host on top, symbiont on bottom) on treeducken test data.

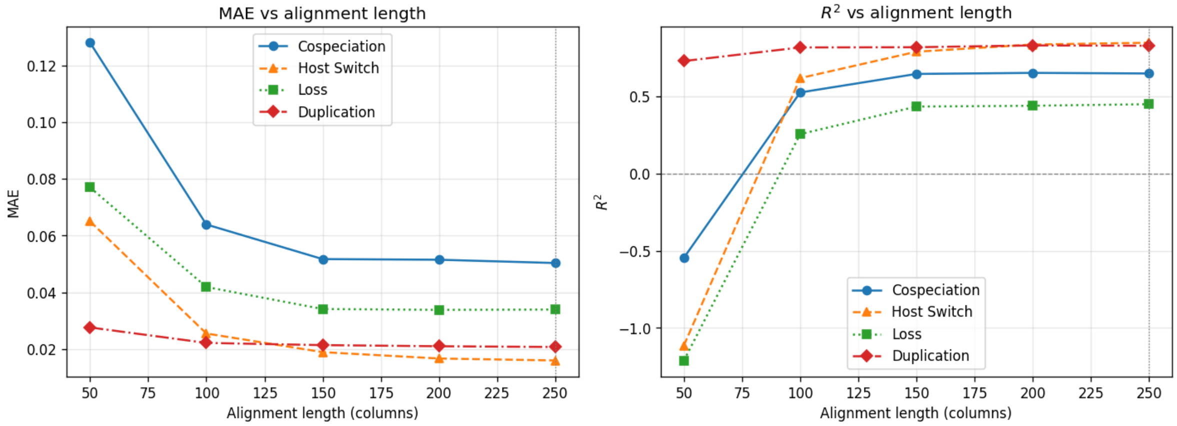


Supplementary Figure 5: Fox accuracy as a function of alignment length. MAE (left) and $R^2$ (right) per event over the 100 held-out AsymmeTree test datasets, as sequences are truncated to 50–250 columns. The dotted line marks the trained 250-column window.

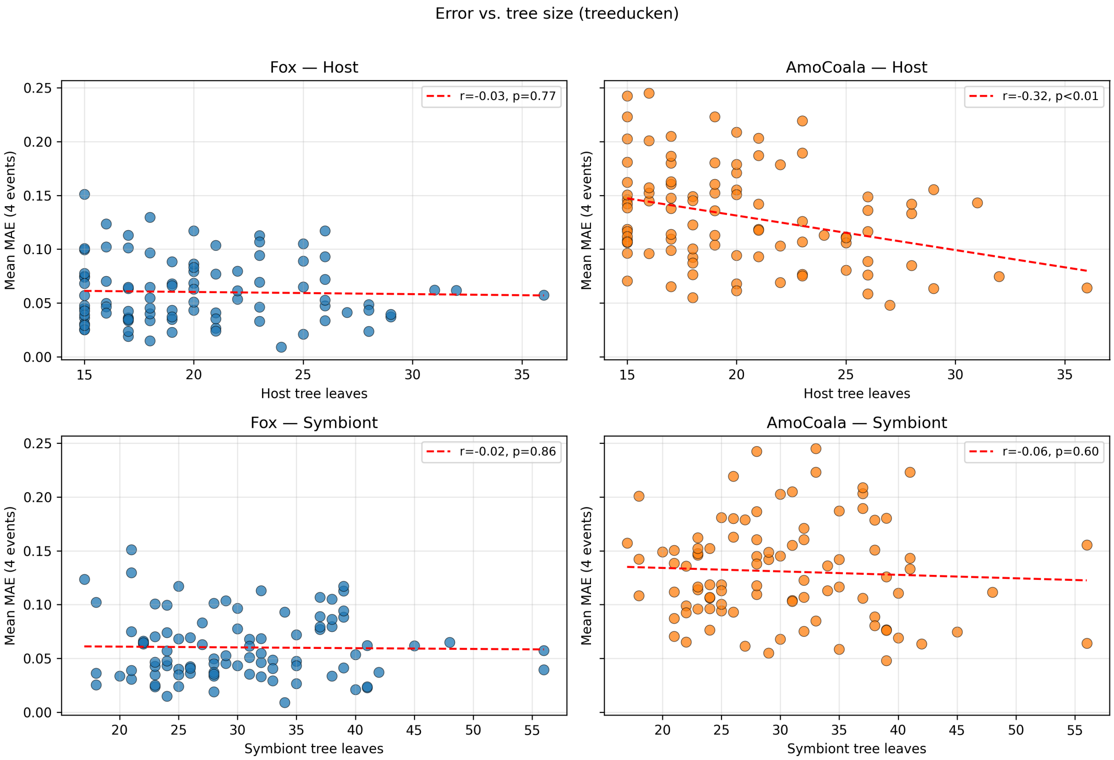


Supplementary Figure 6: AmoCoala vs Fox mean error as a function of tree size on treeducken test data.

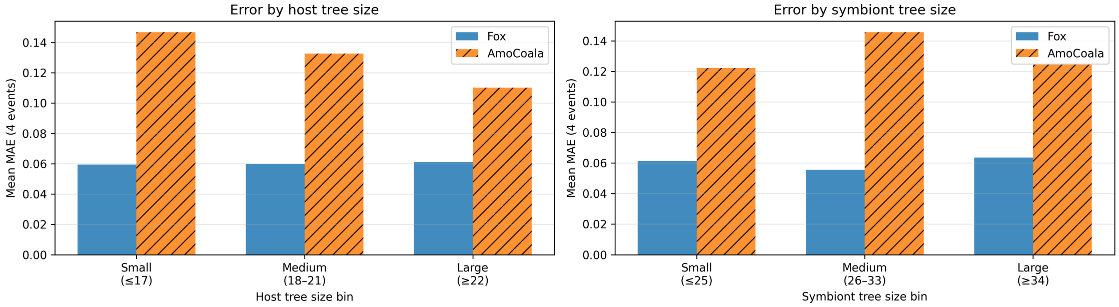


Supplementary Figure 7: AmoCoala vs Fox mean error binned by tree size on treeducken test data. Values are given in Supplementary Table 4.

Supplementary Table 4: Mean absolute error (averaged over the four events) by tree-size tertile, on the 92 TREEDUCKEN test datasets completed by AMOCOALA (3 rounds). Symbiont leaves include lost lineages. FOX stays between 0.056 and 0.064 in every group; AMOCOALA is about two to two and a half times higher in every group. The comparison excludes the 8 largest datasets, on which AMOCOALA did not finish.

| Tree | Tertile (leaves) | $n$ | FOX MAE | AMOCOALA MAE |
|---|---|---|---|---|
| Host | Small ($\leq 17$) | 34 | 0.0595 | 0.1467 |
| | Medium (18–21) | 29 | 0.0599 | 0.1327 |
| | Large ($\geq 22$) | 29 | 0.0612 | 0.1103 |
| Symbiont | Small ($\leq 25$) | 32 | 0.0615 | 0.1221 |
| | Medium (26–33) | 31 | 0.0556 | 0.1456 |
| | Large ($\geq 34$) | 29 | 0.0636 | 0.1246 |

Supplementary Table 5: Invariance of FOX to the order of sequences in the MSAs. For each of the 100 ASYMMETREE held-out test datasets, FOX was run on the original order and on 5 random permutations of the host rows and of the symbiont rows (distance matrices permuted accordingly, associations rebuilt from leaf names; 600 forward passes, no identity permutation). $\Delta$ is the permuted prediction minus the original one; the prediction range is max minus min of the original predictions across datasets. All deviations are at float32 round-off level ($\leq 2.4 \times 10^{-7}$).

| Event | max $\|\Delta\|$ | mean $\|\Delta\|$ | prediction range | max $\|\Delta\|$ / range |
|---|---|---|---|---|
| Cospeciation | $2.38 \times 10^{-7}$ | $2.70 \times 10^{-8}$ | 0.482 | $4.9 \times 10^{-7}$ |
| Host-switch | $5.96 \times 10^{-8}$ | $1.08 \times 10^{-8}$ | 0.218 | $2.7 \times 10^{-7}$ |
| Loss | $2.01 \times 10^{-7}$ | $1.68 \times 10^{-8}$ | 0.201 | $1.0 \times 10^{-6}$ |
| Duplication | $7.45 \times 10^{-8}$ | $1.79 \times 10^{-8}$ | 0.331 | $2.2 \times 10^{-7}$ |

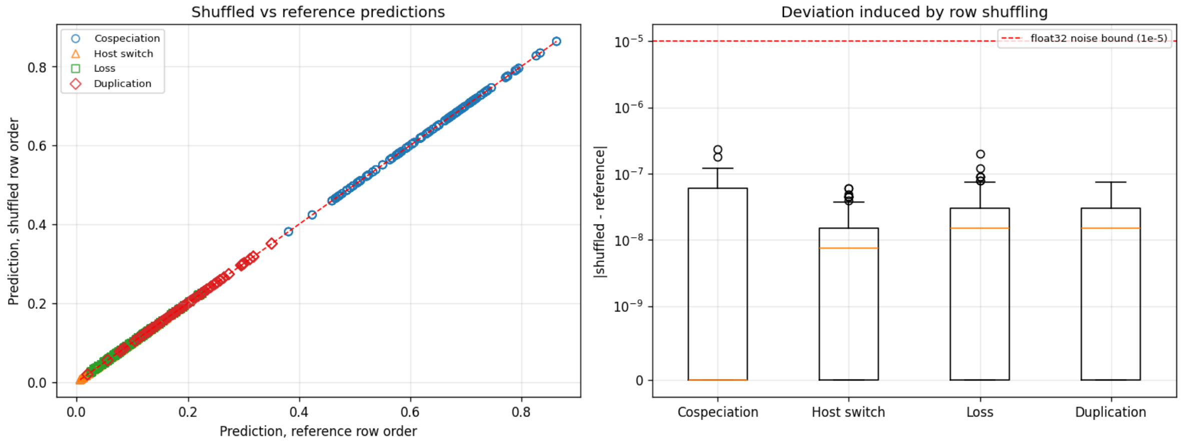


Supplementary Figure 8: Invariance of FOX to sequence order (see Supplementary Table 5). Left: predictions on row-shuffled inputs against predictions on the original order, all events. Right: absolute deviation per event (log scale); the dashed line marks a $10^{-5}$ float32 noise bound.

Supplementary Table 6: Fox on the cockroach–*Blattabacterium* data (52 pairs) as a function of the number of alignment columns read. Each alignment is read up to its own length (the host alignment has 3,736 columns). Run time on CPU (Apple M4 Pro); reading the complete alignments used about 16 GB of memory.

| Columns | Time | Cosp. | Host switch | Loss | Dup. |
|---|---|---|---|---|---|
| 250 | 0.3 s | 0.610 | 0.169 | 0.146 | 0.076 |
| 500 | 0.7 s | 0.676 | 0.168 | 0.116 | 0.039 |
| 1,000 | 1.8 s | 0.767 | 0.127 | 0.075 | 0.031 |
| 2,000 | 5.0 s | 0.674 | 0.196 | 0.101 | 0.028 |
| 4,000 | 15 s | 0.663 | 0.179 | 0.121 | 0.036 |
| 8,000 | 36 s | 0.670 | 0.166 | 0.126 | 0.039 |
| 16,000 | 113 s | 0.684 | 0.163 | 0.117 | 0.036 |
| Full (35,310) | 6–10 min | 0.705 | 0.157 | 0.106 | 0.033 |
